# Noise control is a primary function of microRNAs and post-transcriptional regulation

**DOI:** 10.1101/168641

**Authors:** Jörn M. Schmiedel, Debora S. Marks, Ben Lehner, Nils Blüthgen

**Affiliations:** Institute of Pathology, Charité – Universitätsmedizin Berlin, 10117 Berlin, Germany; IRI Life Sciences, Humboldt Universität Berlin, 10115 Berlin, Germany; EMBL-CRG Systems Biology Unit, Centre for Genomic Regulation (CRG), European Molecular Biology Organization, 08003 Barcelona, Spain; Universitat Pompeu Fabra, 08003 Barcelona, Spain; Department of Systems Biology, Harvard Medical School, Boston, MA 02115, USA; Institució Catalana de Recerca i Estudis Avançats, 08010 Barcelona, Spain; Berlin Institute of Health, 10178 Berlin, Germany

## Abstract

microRNAs are pervasive post-transcriptional regulators of protein-coding genes in multicellular organisms. Two fundamentally different models have been proposed for the function of microRNAs in gene regulation. In the first model, microRNAs act as repressors, reducing protein concentrations by accelerating mRNA decay and inhibiting translation. In the second model, in contrast, the role of microRNAs is not to reduce protein concentrations *per se* but to reduce fluctuations in these concentrations. Here we present genome-wide evidence that mammalian microRNAs frequently function as noise controllers rather than repressors. Moreover, we show that post-transcriptional noise control has been widely adopted across species from bacteria to animals, with microRNAs specifically employed to reduce noise in regulatory and context-specific processes in animals. Our results substantiate the detrimental nature of expression noise, reveal a universal strategy to control it, and suggest that microRNAs represent an evolutionary innovation for adaptive noise control in animals.

**Highlights:**

- Genome-wide evidence that microRNAs function as noise controllers for genes with context-specific functions
- Post-transcriptional noise control is universal from bacteria to animals
- Animals have evolved noise control for regulatory and context-specific processes

## Introduction

MicroRNAs are ubiquitous post-transcriptional repressors of protein-coding genes in multi-cellular organisms (Lee et al., 1993; Reinhart et al., 2000; Reinhart et al., 2002; Wightman et al., 1993). Animal microRNAs bind to ~7nt short, complimentary sequence motifs in the 3’ untranslated regions (3’UTR) of genes (Enright et al., 2003; Lewis et al., 2003; Stark et al., 2003) and, together with cofactors (Hutvágner and Zamore, 2002), accelerate mRNA decay and inhibit translation (Lee et al., 1993; Lim et al., 2005; Wightman et al., 1993). The repertoire of animal microRNA genes has undergone repeated bursts of expansion in various evolutionary lineages (Berezikov, 2011; Fromm et al., 2015; Grimson et al., 2008; Hertel et al., 2006), with around 500 bona fide human microRNAs (Fromm et al., 2015). As many as 60% of human protein coding genes are estimated to have evolutionary constrained microRNA target sites (Friedman et al., 2009). Among the preferential targets of microRNAs are transcription factors and developmental genes (Enright et al., 2003; John et al., 2004; Lewis et al., 2003; Stark et al., 2005). In contrast, highly expressed genes with housekeeping functions are depleted of microRNA binding sites and have thus been dubbed anti-targets (Farh et al., 2005; Stark et al., 2005).

Two fundamentally different roles have been proposed for the function of microRNAs in gene regulation. The first proposed role is the repression of protein output, which has emerged as the dominant paradigm highlighted by the seminal discovery of microRNAs acting as switch-like repressors in *C. elegans* larval development (Lee et al., 1993; Reinhart et al., 2000; Wightman et al., 1993). Subsequent large-scale studies have revealed that microRNAs typically repress tens to hundreds of genes, albeit mostly to a moderate (less than two-fold) degree (Baek et al., 2008; Lim et al., 2005; Selbach et al., 2008). This gave rise to a view of microRNAs as quantitative repressors of protein output that act as a regulatory layer on top of transcriptional regulation (Bartel and Chen, 2004; Shkumatava et al., 2009; Sood et al., 2006; Stark et al., 2005).

In contrast to such a conventional repressive role, microRNAs have also been proposed to regulate genes in order to reduce random fluctuations in their protein concentrations (Bartel and Chen, 2004; Ebert and Sharp, 2012; Hornstein and Shomron, 2006; Osella et al., 2011; Siciliano et al., 2013). Recent work using single cell reporters demonstrated that microRNAs reduce expression noise, i.e. exert ‘noise control’, if microRNA-mediated repression is offset by a compensatory increase in transcription (Schmiedel et al., 2015). Indicative of ‘noise control’, microRNAs have been found to reduce phenotypic variability during animal development (Hilgers et al., 2010; Kasper et al., 2017; Kim et al., 2013; Li et al., 2009; Li et al., 2006; Medeiros et al., 2011).

Gene expression noise is an inevitable consequence of the stochasticity of chemical reactions involved in protein production (Elowitz et al., 2002; McAdams and Arkin, 1997; Thattai and van Oudenaarden, 2001). Expression noise counteracts the precise control of protein levels (Raj and van Oudenaarden, 2008) and thereby presumably impacts organismal fitness (Fraser et al., 2004; Lehner, 2008; Newman et al., 2006). Support for a detrimental impact of expression noise on fitness comes mostly from yeast, where genes that elicit growth defects upon perturbations to their expression levels (hereafter expression-sensitive genes) have low protein noise (Bar-Even et al., 2006; Batada and Hurst, 2007; Lehner, 2008; Newman et al., 2006). Consistently, expression-sensitive genes in yeast show evidence of regulatory patterns thought to minimize expression noise, namely clustering in regions of open chromatin (Batada and Hurst, 2007), avoidance of noisy promoter architectures (Keren et al., 2016; Lehner, 2010; Weinberger et al., 2012) and increased usage of mRNA to translate proteins (Fraser et al., 2004). In multicellular animals, however, there is little evidence for whether or how expression noise is regulated.

Here we explore the relative dominance of the two proposed roles of microRNAs by investigating the relationship between microRNA regulation and transcriptional regulation of genes. If microRNAs act as repressors of protein output we would expect microRNA-mediated repression to either positively correlate with transcriptional *repression* of target genes (for switch-like interactions) or show no correlation (for quantitative repression). In contrast, for microRNAs to act as ‘noise controllers’, their repressive effects must be compensated by transcriptional *activation* of target genes.

To discriminate between the two models of microRNA function we investigate transcriptional and post-transcriptional expression parameters of genes in a mouse cell line and find that the majority of microRNA targets exhibit compensatory transcription. Further substantiating a role for microRNAs in noise control, target set analysis shows that microRNAs preferentially regulate expression-sensitive disease and developmental genes in human and mouse. Noise control is, however, not unique to microRNAs, and an analysis of transcriptome and proteome data shows that post-transcriptional noise control is a general strategy in organisms ranging from bacteria to animals. Finally, a comparative analysis of noise control patterns across eukaryotes reveals animal-specific gains of noise control in regulatory and context-specific processes. These gains strongly correlate with the enrichment of microRNA target sites, suggesting microRNAs have been instrumental in shaping noise control in animals.

## Results

### MicroRNA target genes have elevated transcription rates

Two fundamentally different models have been proposed for microRNAs in gene regulation. In the first model, microRNAs act as simple repressors, to switch-off (Lee et al., 1993; Reinhart et al., 2000; Wightman et al., 1993) or quantitatively reduce protein concentration (Bartel and Chen, 2004; Shkumatava et al., 2009; Sood et al., 2006) of target genes. In the second model, the primary function of microRNAs is not to reduce protein concentrations *per se*, but rather to stabilize expression, i.e. to control gene expression noise (Bartel and Chen, 2004; Ebert and Sharp, 2012; Osella et al., 2011; Schmiedel et al., 2015; Siciliano et al., 2013).

We reasoned that investigating the transcription rates of microRNA target genes would allow us to distinguish between these two alternative hypotheses. If microRNAs were to repress protein output, their targets should be genes that already have low transcription rates. On the other hand, if microRNAs were to reduce expression variability, their target genes should display compensatory transcriptional activation that offsets the reduction in average protein output (Figure 1A) (Ebert and Sharp, 2012; Herranz and Cohen, 2010; Schmiedel et al., 2015).

**Figure 1.**
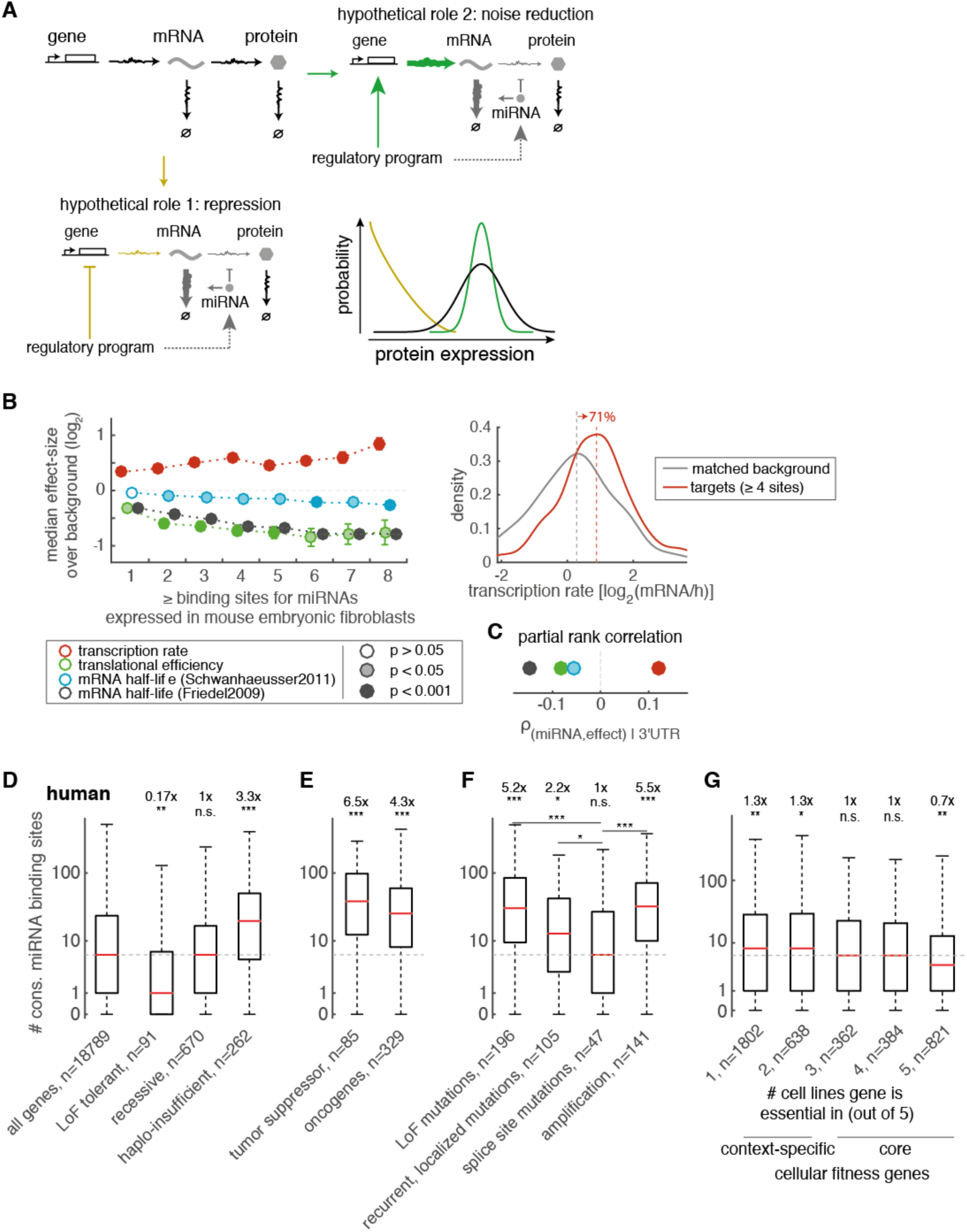
– Noise control is a primary function of human and mouse microRNAs. (A) Schematic of gene expression with possible outcomes of different coordination between a gene’s transcriptional and microRNA regulation. If a regulatory program inhibits a gene’s transcription but promotes its regulation via microRNAs, protein expression is repressed (yellow). On the contrary, if a regulatory program activates a gene’s transcription as well as its regulation via microRNAs, protein expression noise is reduced (green). (B) Coordination of microRNA regulation and transcription in mouse embryonic fibroblasts (MEF). Left panel: Median difference (log_2_) in expression parameters between microRNA targets (with equal or greater than a certain number of conserved binding sites for MEF-expressed microRNAs) and 3’UTR length matched background sets of genes lacking microRNA binding sites. Circles show median differences and error bars the standard deviation of differences across 1000 iterations of background sampling are shown. Fill of circles indicates mean p-value across iterations. empty: p > 0.05, shaded: p < 0.05, filled: p < 0.001 (two-sided Wilcoxon rank sum test). Right panel: Example showing transcription rate distributions (kernel density estimates) of microRNA targets with equal or greater than four sites (red) and their matched background control set (grey). Vertical dashed lines indicate medians. 71% of target genes have transcription rates greater than the median of the background. n = 301. (C) Spearman partial rank correlation coefficients between conserved binding sites for microRNAs expressed in MEF and expression parameters, while controlling for 3’UTR length of genes. Circles are colored and filled as in (B). (D-G) Distributions of conserved microRNA binding sites in 3’UTRs of human (D) genes with varying expression-sensitivity, (E) cancer driver genes, (F) genes enriched for specific alterations in cancer and (G) cellular fitness genes, stratified by the number of cell lines they are essential in. Red bar indicates median, boxes the IQR, whiskers extend to 1.5∗IQR from lower or upper quartile. Outliers are not shown. Horizontal dashed grey line indicates median number of conserved microRNA binding sites across all human genes (first column in D). Indicated above each column are the relative difference in median numbers of microRNA binding sites between genes in the set and genes not in the set as well as the significance level of the difference. In (F and G) the significance of differences between sets are also indicated. LoF: loss-of-function. ^∗^: p < 0.05, ^∗∗^: p < 10^−3^, ^∗∗∗^: p < 10^−6^ (two-sided Wilcoxon rank sum test)

To this end we re-analyzed gene expression parameters in mouse embryonic fibroblasts (MEF) (Friedel et al., 2009; Schwanhäusser et al., 2011). To investigate the dependency of expression parameters on microRNA regulation, we compared genes that contain conserved binding sites for MEF-expressed microRNAs to control sets of genes with matched 3’UTR length but no conserved binding sites. As expected, mRNA half-lives and translational efficiencies of genes decrease with the number of microRNA sites. Importantly, we also find that transcription rates of genes increase with the number of microRNA sites (Figure 1B). Moreover, the effect size of compensatory transcription is equivalent to that of the repressive post-transcriptional effects (up to 1.8-fold increase in transcription rates, up to 1.8-fold, 1.2-fold and 1.7-fold decrease in translation rate constants and mRNA half-lives (from (Schwanhäusser et al., 2011) and (Friedel et al., 2009)), respectively; p < 10^−3^, except mRNA half-lives from (Schwanhäusser et al., 2011): p < 0.05, Wilcoxon rank sum test). These results therefore suggest that a majority of microRNA-target interactions serve to control noise rather than repress protein output.

Equivalent conclusions are reached when examining the partial correlation between the expression parameters and the number of microRNA sites, while controlling for 3’UTR length (Spearman partial rank correlation, R = 0.12, p < 10^−6^, for transcription rates; R = -0.08, p < 10^−3^, for translational rate constants; R = -0.06, p < 0.01, for mRNA half-lives from (Schwanhäusser et al., 2011); R = -0.15, p < 10^−6^, for mRNA half-lives from (Friedel et al., 2009), Figure 1C). In contrast, we do not find increased transcription rates for genes regulated by two other repressive post-transcriptional mechanisms (Supplementary Figure 1), suggesting that compensatory transcription is not *per se* associated with posttranscriptional repression. Of note, lowly expressed genes (lowest ~25%) are underrepresented in the underlying mass spectrometry based dataset (Schwanhäusser et al., 2011), and results therefore do not necessarily hold for this subset of genes. Nevertheless, the data suggest that microRNAs in MEF primarily act to reduce noise of robustly expressed target genes.

### MicroRNAs preferentially target expression sensitive genes

If a primary role of microRNAs is to reduce noise of their target genes, then expression-sensitive genes, for which gene expression noise is most detrimental (Bar-Even et al., 2006; Fraser et al., 2004; Lehner, 2008; Newman et al., 2006), should be preferential microRNA targets. We collected several sets of known human expression-sensitive and expression-insensitive genes and analyzed if expression-sensitivity is associated with the number of conserved microRNA binding sites in 3’UTRs. Haplo-insufficient disease genes (Dang et al., 2008) represent a group of genes where a 50% reduction in gene dosage results in a strong phenotype, i.e. these genes are expression-sensitive. We find that this group of genes is significantly enriched for microRNA binding sites when compared to the remainder of human genes (3.3-fold increase, p < 10^−6^, Fisher’s exact test, Figure 1D). In contrast, genes for which loss of function is tolerated are significantly depleted (5.9-fold decrease, p < 10^−3^, Fisher’s exact test) and genes with recessive disease phenotypes are neither enriched nor depleted of microRNA binding sites (MacArthur et al., 2012).

Equivalently, oncogenes and tumor suppressors (Davoli et al., 2013; Futreal et al., 2004) are also enriched for microRNA binding sites (6.5-fold increase, p < 10^−6^, for tumor suppressors; 4.3-fold increase, p < 10^−6^, for oncogenes, Fisher’s exact test, Figure 1E). While tumor suppressors largely function in an expression-dependent way, oncogenes contribute through both expression-dependent and -independent mechanisms to tumorigenesis. We thus classified cancer genes as expression-dependent and -independent using data from a meta-analysis of tumor sequencing data (Davoli et al., 2013). We find that genes significantly enriched for expression-altering mutations typically associated with tumor suppressors – mutations causing loss-of-function or affecting splice sites – are significantly enriched for microRNA binding sites (5.2-fold increase, p < 10^−6^, for genes enriched in loss-of-function mutations; 2.2-fold increase, p < 0.05, for genes enriched in splice site mutations, Fisher’s exact test, Figure 1F). Similarly, oncogenes enriched for copy-number amplifications are enriched for microRNA binding sites (5.5-fold increase, p < 10^−6^, Fisher’s exact test). On the contrary, oncogenes commonly acting through expression-independent mechanisms in cancer – those with recurrent, localized mutations - are not enriched for microRNA binding sites (Figure 1F). Moreover, genes acting through expression-dependent mechanisms are significantly enriched compared to genes acting through expression-independent mechanisms in cancer (p < 10^−6^, p < 0.05 and p < 10^−6^ for genes enriched in loss-of-function mutations, splice site mutations or copy-number amplifications compared to genes enriched in recurrent, localized mutations, Wilcoxon rank sum test).

Similar to these findings in human, we find mouse genes that are essential for embryonic development (Eppig et al., 2015; Georgi et al., 2013) to be significantly enriched for conserved microRNA binding sites compared to nonessential genes (2-fold increase, p < 10^−6^, Wilcoxon rank sum test, Supplementary Figure 2).

Genes essential for cell growth (hereafter ‘cellular fitness genes’) are another set of genes expected to be in need of noise control. In yeast, cellular fitness genes have low expression noise (Newman et al., 2006) and, presumably, this is in part due to a combination of high transcription rates and low translation rates (Fraser et al., 2004). However, analyzing a set of human cellular fitness genes (Hart et al., 2015) – which is strongly enriched for genes with orthologues in yeast – we find that they are hardly enriched for microRNA binding sites (1.3-fold increase, p < 10^−3^ and p < 0.05, for genes essential in one or two out of five cell lines, respectively; Fisher’s exact test, Figure 1G); and cellular fitness genes that are essential in all five tested cell lines are significantly depleted of microRNA binding sites (1.4-fold decrease, p < 10^−3^, Fisher’s exact test). This therefore suggests either that human cellular fitness genes are - contrary to yeast - not under posttranscriptional noise control, or that such control is exerted by mechanisms other than microRNAs.

Taken together, these results show that, for the subset of developmental and disease genes in human and mouse, expression-sensitivity is strongly linked to microRNA targeting, suggesting microRNAs control noise for these expression-sensitive genes.

### mRNA/protein ratios control gene expression noise

To elucidate why human cellular fitness genes are not microRNA-regulated, we sought a more general approach to study post-transcriptional noise control. Posttranscriptional noise control mechanisms that act (partially) through translational inhibition result in the use of higher mRNA levels to produce a certain amount of protein (Figure 2A) (Blake et al., 2003; Fraser et al., 2004; Ozbudak et al., 2002; Raser and O’Shea, 2004; Schmiedel et al., 2015; Thattai and van Oudenaarden, 2001).

**Figure 2.**
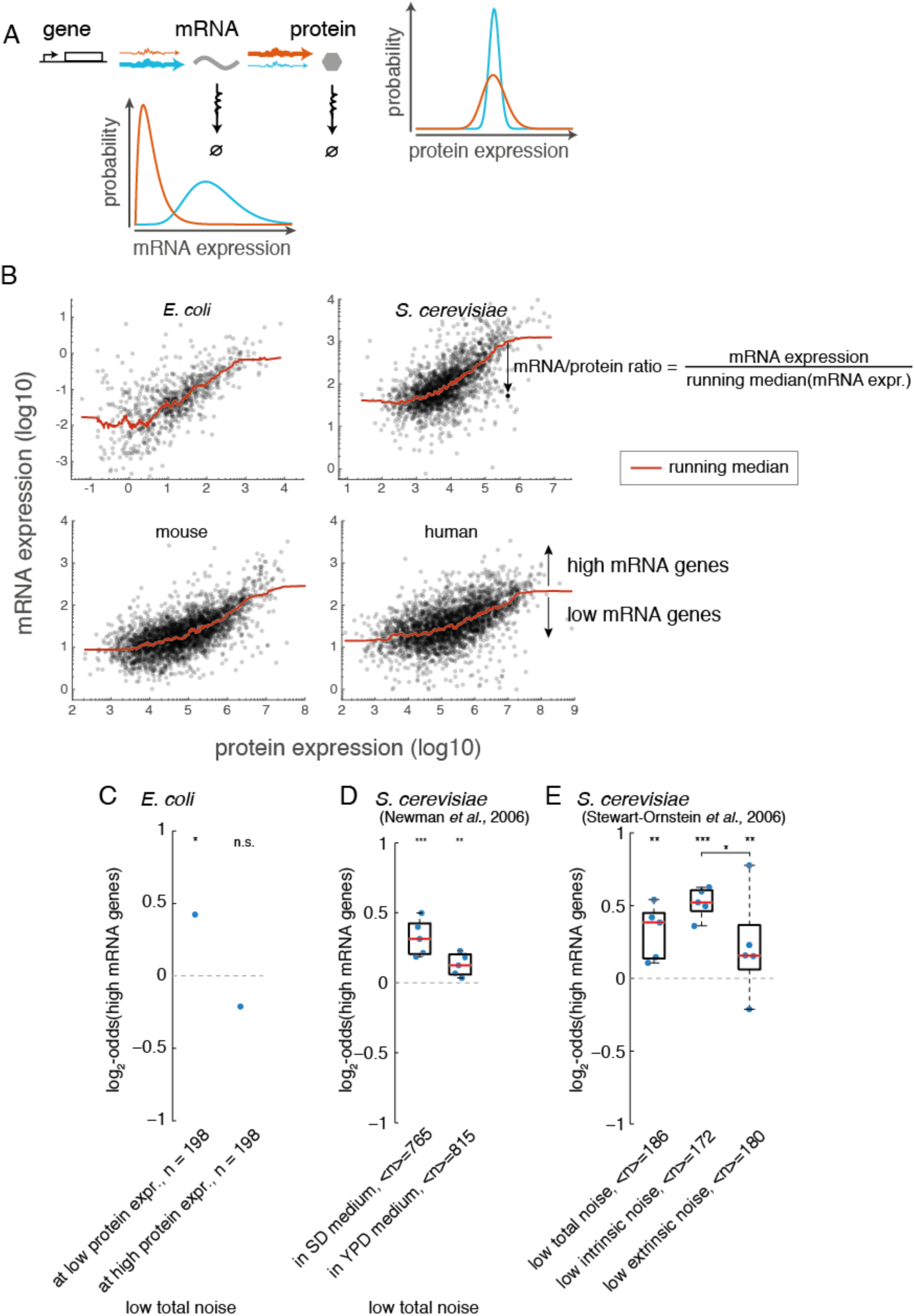
– mRNA/protein ratios control gene expression noise. (A) Scheme of gene expression showing two opposing strategies to express a certain average amount of protein. Low transcription rates but high translation rates will result in low mRNA levels and high protein expression noise (orange). High transcription rates but low translation rates will result in high mRNA levels and low protein expression noise (blue). (B) Genome-wide mRNA and protein expression patterns in *E. coli*, *S. cerevisiae*, mouse (embryonic fibroblasts) and human (kidney). Each dot represents a gene. Red line gives running median of mRNA expression across protein expression range. mRNA/protein ratios are the relative deviation from the running median. Additionally, genes above running median are classified as *high mRNA genes*, others as *low mRNA genes*. (C) Enrichment of *high mRNA genes* among genes with low noise in *E.coli.* Genes are split into low (below median) and high (above median) protein expression (also see Figure S2). (D) Enrichment of high mRNA genes among genes with low total noise in haploid yeast (Newman et al., 2006) in two different growth conditions. SD: synthetic defined, YPD: yeast extract peptone dextrose. Each dot represents a matched transcriptome and proteome datasets. (E) Same as (D) but for diploid yeast, where total noise was also deconvoluted into intrinsic and extrinsic noise components (Stewart-Ornstein et al., 2012). <*n*> denotes the mean number of genes in a set across all pairs of datasets (D and E). Fisher’s Exact test for enrichment (C-E). Wilcoxon rank sum test for differential enrichment between gene sets (E). p-values are aggregated using Fisher’s Method. ^∗^: p < 0.05, ^∗∗^: p < 10^−3^, ^∗∗∗^: p < 10^−6^.

We thus searched for patterns of such altered mRNA/protein ratios using transcriptome and proteome expression data and investigated their relation to gene expression noise and microRNA regulation. To this end we collected quantitative transcriptomes and proteomes for *E. coli* (Taniguchi et al., 2010), *S. cerevisiae* (Kulak et al., 2014; Lawless et al., 2016; Presnyak et al., 2015), *Capsaspora owczarzaki* (Sebé-Pedrós et al., 2013; Sebé-Pedrós et al., 2016b), mouse (Geiger et al., 2013; Jovanovic et al., 2015; Merkin et al., 2012; Schwanhäusser et al., 2011) and human (Khan et al., 2013; Wilhelm et al., 2014) (see Methods). In agreement with previous reports (Lawless et al., 2016; Schwanhäusser et al., 2011; Sebé-Pedrós et al., 2016b; Vogel et al., 2010; Wilhelm et al., 2014), transcriptome and proteome are moderately correlated and, consistently across organisms, proteins with similar levels are associated with mRNA levels that can differ by up to two orders of magnitude (Figure 2B).

For each gene in a matched transcriptome/proteome dataset, we defined the mRNA/protein ratio as the relative deviation of its mRNA levels from the running median of mRNA levels across the protein expression range. We additionally classified genes in a binary fashion as “high mRNA genes” or “low mRNA genes”, depending on whether their mRNA levels are above or below the running median at their given protein expression (Figure 2B). This approach allows normalizing for general trends as well as for varying magnitudes of variability of mRNA levels along the protein expression range. We find that mRNA/protein ratios are well suited to investigate differential translation rates of genes (Supplementary Figure 3) as well as the impact of post-transcriptional regulatory mechanisms (Supplementary Figure 4).

We first set out to test whether larger mRNA/protein ratios are associated with lower protein expression variability genome-wide, as predicted previously (Fraser et al., 2004; Ozbudak et al., 2002; Raser and O’Shea, 2004). To this end we used genome-wide measurements of protein expression noise in *E. coli* and budding yeast. We calculated noise/protein ratios along the protein expression range equivalently to mRNA/protein ratios and classified genes as ‘low noise genes’ and ‘high noise genes’ (Figure S2).

For genes in *E. coli* we calculated mRNA/protein ratios and noise/protein ratios based on measurements from RNA-sequencing and YFP-fusion proteins (Taniguchi et al., 2010) (Supplementary Figure 5). We find a significant enrichment of high mRNA genes among low noise genes in the lower half of the protein expression range (log_2_-odds ratio (OR) = 0.42, p < 10^−3^, Fisher’s exact test, Figure 2D). On the contrary, in the upper half of the protein expression range we could not detect a significant enrichment (OR = -0.21, p = 0.6, Fisher’s exact test). This observation is consistent with the notion that, in *E. coli*, noise of highly expressed proteins is dominated by extrinsic components, which should not be affected by mRNA/protein ratios, whereas noise of lowly expressed proteins is dominated by intrinsic noise, that is noise from the production process of proteins (Taniguchi et al., 2010). Therefore, we conclude that at low expression levels, i.e. where intrinsic noise matters, high mRNA/protein ratios are associated with low protein expression variability in *E. coli*.

For budding yeast, we calculated mRNA/protein ratios from RNA-sequencing and mass spectrometry data (Kulak et al., 2014; Lawless et al., 2016; Presnyak et al., 2015) and noise/protein ratios from GFP-fusion protein data (Newman et al., 2006; Stewart-Ornstein et al., 2012) (see Methods). We find a significant enrichment of high mRNA genes among genes with low total protein expression noise (median(OR) = 0.31, p < 10^−6^, for total noise in SD medium; median(OR) = 0.12, p < 10^−3^, for total noise in YPD medium; median(OR) = 0.38, p < 10^−3^, for total noise in diploid strains; Fisher’s exact test, Figure 2E&F). A decomposition of total noise into intrinsic and extrinsic noise components, based on the comparison of diploid strains tagged with one or two fluorescent proteins (Stewart-Ornstein et al., 2012), further shows that high mRNA genes are more enriched among genes with low intrinsic noise than low extrinsic noise (difference in median(OR) = 0.37, p < 0.05, Wilcoxon rank sum test, Figure 2F), consistent with high mRNA/protein ratios leading to low intrinsic protein expression noise also in budding yeast.

Genome-wide expression and noise data from *E. coli* and budding yeast therefore validate established biophysical models (Blake et al., 2003; Ozbudak et al., 2002; Raser and O’Shea, 2004; Thattai and van Oudenaarden, 2001) in that - at equal protein expression - high mRNA levels promote low intrinsic noise.

### Expression-sensitive genes are enriched for large mRNA/protein ratios across species

If mRNA/protein ratios were indeed signatures of post-transcriptional noise control, then expression-sensitive genes should have large mRNA/protein ratios. In *E. coli*, essential genes - as defined by non-viability upon knockout (Hashimoto et al., 2005) – are enriched among high mRNA genes as well as low noise genes (OR = 0.48, p < 0.05 and OR = 1.11, p < 10^−3^, respectively; Fisher’s exact test, Figure 3 A&B). In yeast, genes with growth phenotypes upon heterozygous deletion (Deutschbauer et al., 2005) or over-expression (Makanae et al., 2013) show an enrichment in high mRNA genes (median(OR) = 0.63, p < 10^−6^ and median(OR) = 0.60, p < 10^−3^, respectively; Fisher’s exact test, Figure 3C), consistent with previous findings for essential genes and protein sub-complexes (Fraser et al., 2004). Consistently, these expression-sensitive genes also display lower intrinsic noise (OR = 0.88, p < 0.05 and OR = 0.65, p = 0.12, respectively; Fisher’s exact test, Figure 3D), similar to results from previous analyses that considered total noise levels (Lehner, 2008; Newman et al., 2006).

**Figure 3.**
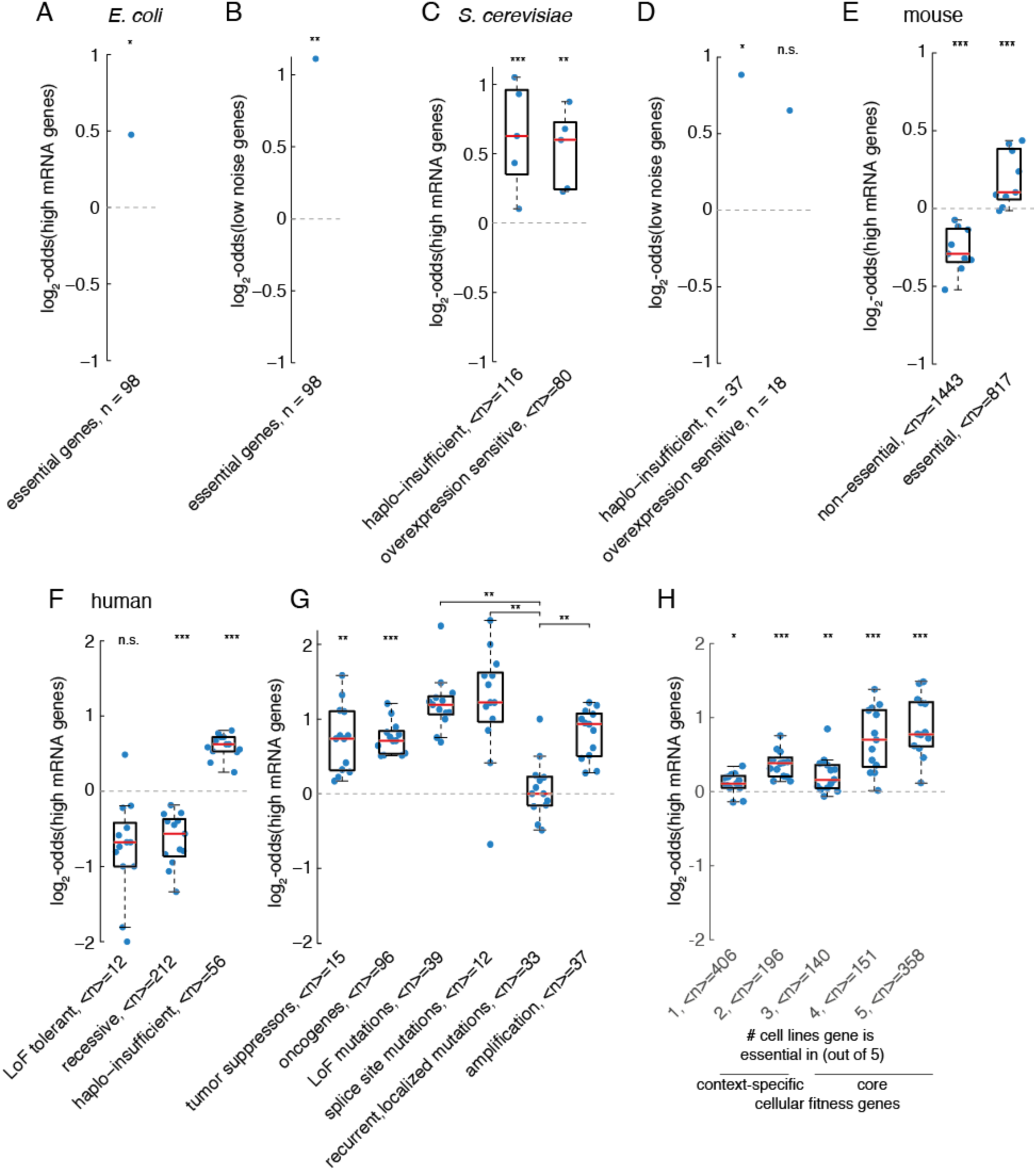
– Expression-sensitive genes are enriched for large mRNA/protein ratios. Enrichment of *high mRNA genes* among (A) essential genes in *E. coli*, (C) yeast genes sensitive to decreased or heightened expression, (E) mouse essential developmental genes, (F) human genes with varying expression-sensitivity, (G) human cancer genes or (H) human cellular fitness genes. (B, D) Enrichment of *low noise genes* among (B) essential genes in *E. coli* and (D) yeast genes sensitive to decreased or heightened expression. LoF: loss-of-function. ^∗^: p < 0.05, ^∗∗^: p < 10^−3^, ^∗∗∗^: p < 10^−6^ (two-sided Wilcoxon rank sum test, p-values aggregated with Fisher’s method)

Genes that are essential in mouse embryonic development are enriched for high mRNA genes, while non-essential genes are depleted of high mRNA genes (median(OR) = 0.1, p < 10^−6^ and median(OR) = -0.29, p < 10^−6^, respectively; Fisher’s exact test, Figure 3E); which is consistent with our finding that microRNA target sites are enriched in these essential genes (see Supplementary Figure 2). Similarly, loss-of-function tolerant and recessive human genes are depleted of high mRNA genes (median(OR) = -0.7, p = 0.13 and median(OR) = -0.51, p < 10^−6^, respectively; Fisher’s exact test), while haplo-insufficient genes, tumor suppressors and oncogenes are enriched for high mRNA genes (median(OR) = 0.62, p < 10^−6^; median(OR) = 0.76, p < 10^−3^ and median(OR) = 0.73, p < 10^−6^, respectively; Fisher’s exact test, Figure 3F & G). Furthermore, while expression-dependent cancer genes are enriched for high mRNA genes (median(OR) = 1.16 for loss-of-function mut. enriched genes; median(OR) = 1.34 for splice site mut. enriched genes; median(OR) = 0.94 for copy-number amp. enriched genes; all p < 10^−6^; Fisher’s exact test), expression-independent oncogenes are not enriched for high mRNA genes (median(OR) = 0, p = 0.27 for genes enriched with recurrent, localized mutations; Fisher’s exact test, Figure 3G).

Interestingly, human cellular fitness genes, a class of genes that shows no enrichment for microRNA binding sites, are enriched for high mRNA genes, and this enrichment increases with the number of cell lines that genes are essential in (median log2-odds from 0.11 to 0.77, p < 0.05 and p < 10^−6^, for genes essential in one or five out of five cell lines, respectively; Fisher’s exact test, Figure 3H). This suggests that human cellular fitness genes are under post-transcriptional noise control, yet they use other mechanisms then microRNA regulation.

Taken together, these results demonstrate that post-transcriptional noise control of expression-sensitive genes by means of large mRNA/protein ratios is a general feature shared between bacteria, uni- and multi-cellular eukaryotes.

### mRNA/protein ratios reveal functional constraints on microRNA-mediated noise reduction

We split human expression-sensitive genes into two groups, depending on whether they are enriched for microRNA binding sites or not. *Core cellular fitness genes* contain all cellular fitness genes that are essential in three to five out of five tested cell lines (Hart et al., 2015) and are on average depleted of microRNA binding sites. In contrast, *context*-*specific expression*-*sensitive genes* contain cellular fitness genes that are essential in one or two cell lines as well as all expression-sensitive disease genes (haplo-insufficient, tumor suppressors and oncogenes) and are enriched for microRNA binding sites. To investigate why core cellular fitness genes are enriched for high mRNA genes but depleted of microRNA sites, we first quantified the 3’UTR length of genes as a function of their average mRNA/protein ratios across human datasets. Interestingly, we find a general trend for shorter 3’UTRs at larger mRNA/protein ratios (1.6-fold decrease in median 3’UTR length of all genes above scaled log-_10_-mRNA/protein ratio = 1, compared to genes below, p < 10^−6^, one-sided Wilcoxon rank sum test, Figure 4B and C), from which core cellular fitness genes further deviate towards even shorter 3’UTRs (2.6-fold decrease in median 3’UTR length above scaled log_10_-mRNA/protein ratio = 1, compared to non-core cellular fitness genes, p < 10^−3^, one-sided Wilcoxon rank sum test).

**Figure 4.**
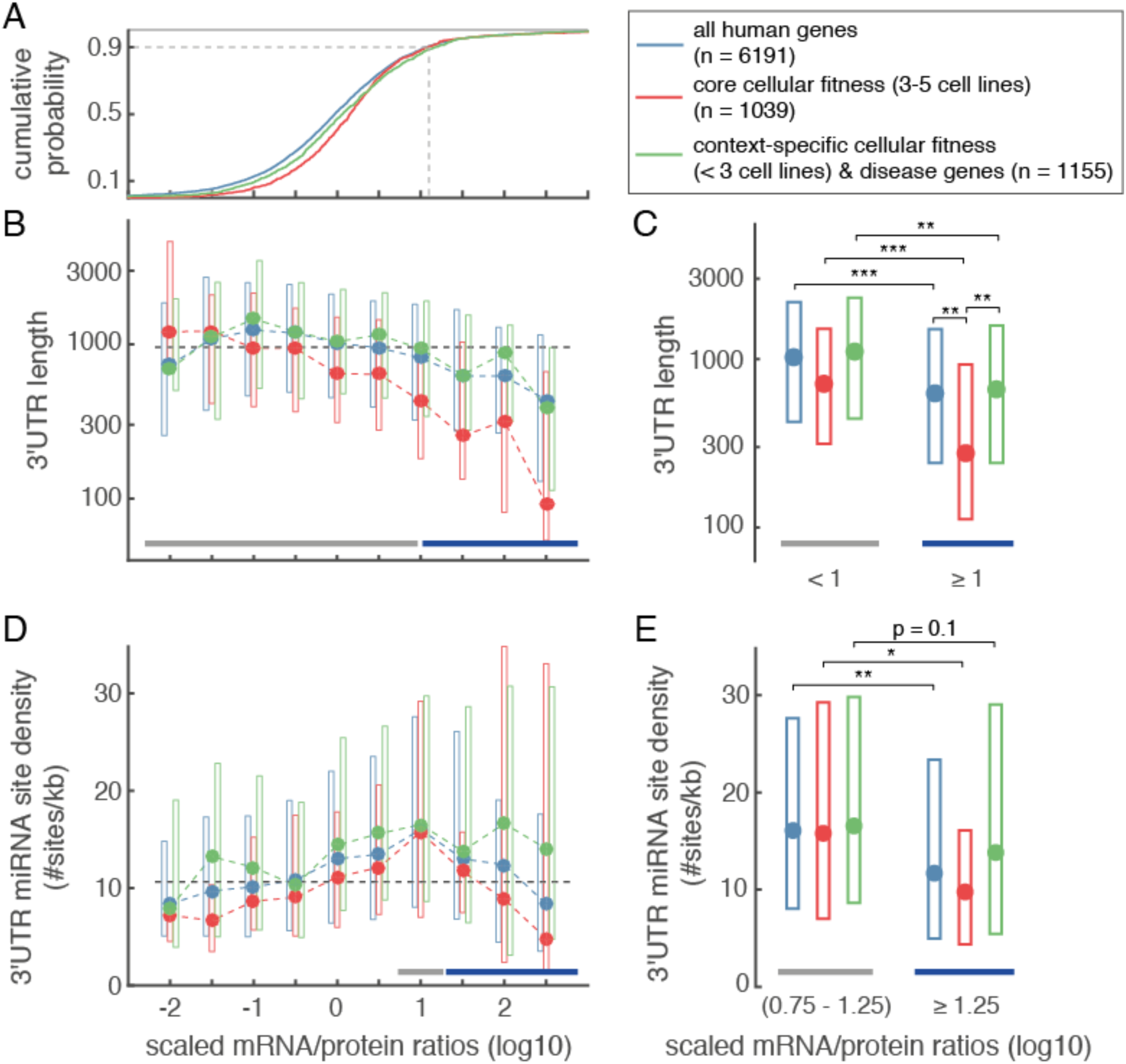
– mRNA/protein ratios reveal functional constrains of microRNA regulation. (A) Cumulative distribution of human genes as function of scaled mRNA/protein ratios (log_10_-transformed, standardized and averaged across datasets, see Methods). For reference, 90^th^ percentile of cumulative distribution is indicated as vertical dashed gray line in scaled mRNA/protein space. (B&D) 3’UTR length (B) and 3’UTR microRNA site density (D) distributions of human genes stratified by mRNA/protein ratios. 3’UTR microRNA site density is the number of conserved microRNA binding sites per kb 3’UTR. Genes were stratified into bins with width 0.5 and centers as indicated. Dots represent medians, boxes the inter-quartile ranges. Black dashed horizontal line indicates median across all human genes considered for analysis. (C) Differences in 3’UTR length between genes with scaled mRNA/protein ratios less than and equal or greater than scaled mRNA/protein ratios = 1, as indicated by grey and blue bars in (B). (E) Differences in 3’UTR microRNA site density between the peak region (scaled mRNA/protein ratios between 0.75 and 1.25) and above (scaled mRNA/protein ratios equal or greater than 1.25), as indicated by grey and blue bars in (D). In (C) and (E): ^∗^: p < 0.05, ^∗∗^: p < 10^−3^ ^∗∗∗^ p < 10^−6^, one-sided Wilcoxon rank sum test.

To assess whether these shorter 3’UTRs reflect an avoidance of microRNA regulation (Farh et al., 2005; Stark et al., 2005), we calculated the density of conserved microRNA sites per kilobase 3’UTR. Stratifying genes by their mRNA/protein ratios shows that microRNA site density increases as a function of mRNA/protein ratios, but drops off again for the 10% of genes with the largest mRNA/protein ratios (1.4-fold decrease in microRNA site density compared to peak region, p < 10^−3^, one-sided Wilcoxon rank sum test, Figure 4 D and E). Core cellular fitness genes show a weak tendency for lower microRNA site density. In contrast, context-specific expression-sensitive genes show a weak tendency for heightened microRNA site density. However, both gene groups show the general trend of a peak and subsequent drop off in microRNA site density (1.6-fold and 1.2-fold decrease compared to peak region, p < 0.05 and p = 0.1, for core cellular fitness and context-specific expression-sensitive genes, respectively, one-sided Wilcoxon rank sum test).

Together, this suggests that core cellular fitness genes are depleted of microRNA sites mainly due to their short 3’UTRs. Because the microRNA site density of core cellular fitness genes is similar to that of other genes, it is unlikely that short 3’UTRs are a consequence of microRNA site avoidance *per se.* Importantly, the analysis suggests that anti-targets, those genes that avoid microRNA regulation (Farh et al., 2005; Stark et al., 2005), are found both at the lower as well as the upper end of the post-transcriptional repression spectrum. Such an avoidance of microRNA sites for the 10% of genes with the largest mRNA/protein ratios (Figure 4A) is in good agreement with the upper limit for microRNA-mediated noise reduction, which we have previously estimated to be above the 90^th^ percentile of mRNA expression (Schmiedel et al., 2015). The microRNA site density profile across mRNA/protein ratios thus resembles what would be expected if it were driven by an increasing need for noise control combined with a limited potential of microRNAs to reduce noise for genes with very large mRNA expression.

The microRNA site density profile across mRNA/protein ratios is therefore consistent with microRNAs acting – within functional constraints – to reduce expression noise.

### Evolution of post-transcriptional noise control in eukaryotes

To investigate how noise control evolved in animals we performed a comparative analysis of mRNA/protein ratios for functional groups of genes. We calculated enrichment of high mRNA genes for Gene Ontology biological process annotations using transcriptome/proteome data sets from budding yeast, mouse and human. To allow for a better assessment of mRNA/protein ratio evolution in animals we also used matched transcriptome/proteome datasets from the three cell types of the amoeboid filasterean *Capsaspora owczarzaki* (here after Capsaspora). Capsaspora is a close unicellular relative to animals, with the ability to temporally differentiate into three distinct cell types, a repertoire of animal-related genes involved in cell differentiation and signaling, but a simpler genomic architecture for transcriptional regulation (Sebé-Pedrós et al., 2016a; Sebé-Pedrós et al., 2013; Sebé-Pedrós et al., 2016b; Suga et al., 2013).

To allow comparison between the four organisms, we restricted our analysis to 34 biological processes that have on average at least 30 annotated genes in the datasets of each organism (see Methods). We also incorporated the enrichment of microRNA-binding sites among genes annotated in these processes in human and mouse into our analysis.

Hierarchical clustering of enrichment patterns across datasets reveals a hierarchy that resembles the phylogeny of organisms. Yeast datasets are distant from the (pre-)animal part of the tree (Figure 5A). Capsaspora datasets are distinct from most animal datasets except for mouse metabolic tissues (see also Supplementary Figure 6A). In mouse and human, data sets with similar functions cluster together, such as metabolic tissues, hormone-producing tissues, tissues related to immune system functions or reproductive tissues. Together this suggests that mRNA/protein ratios are functionally meaningful and subject to evolutionary forces, which is further supported by significant gene-level correlations of mRNA/protein ratios between organisms (Spearman rank correlation, R = 0.19, p < 10^−6^ for human-yeast orthologues; R = 0.25, p < 10^−6^ for human-Capsaspora orthologues; R = 0.5, p < 10^−6^ for human-mouse orthologues; Supplementary Figure 7 A-C).

**Figure 5.**
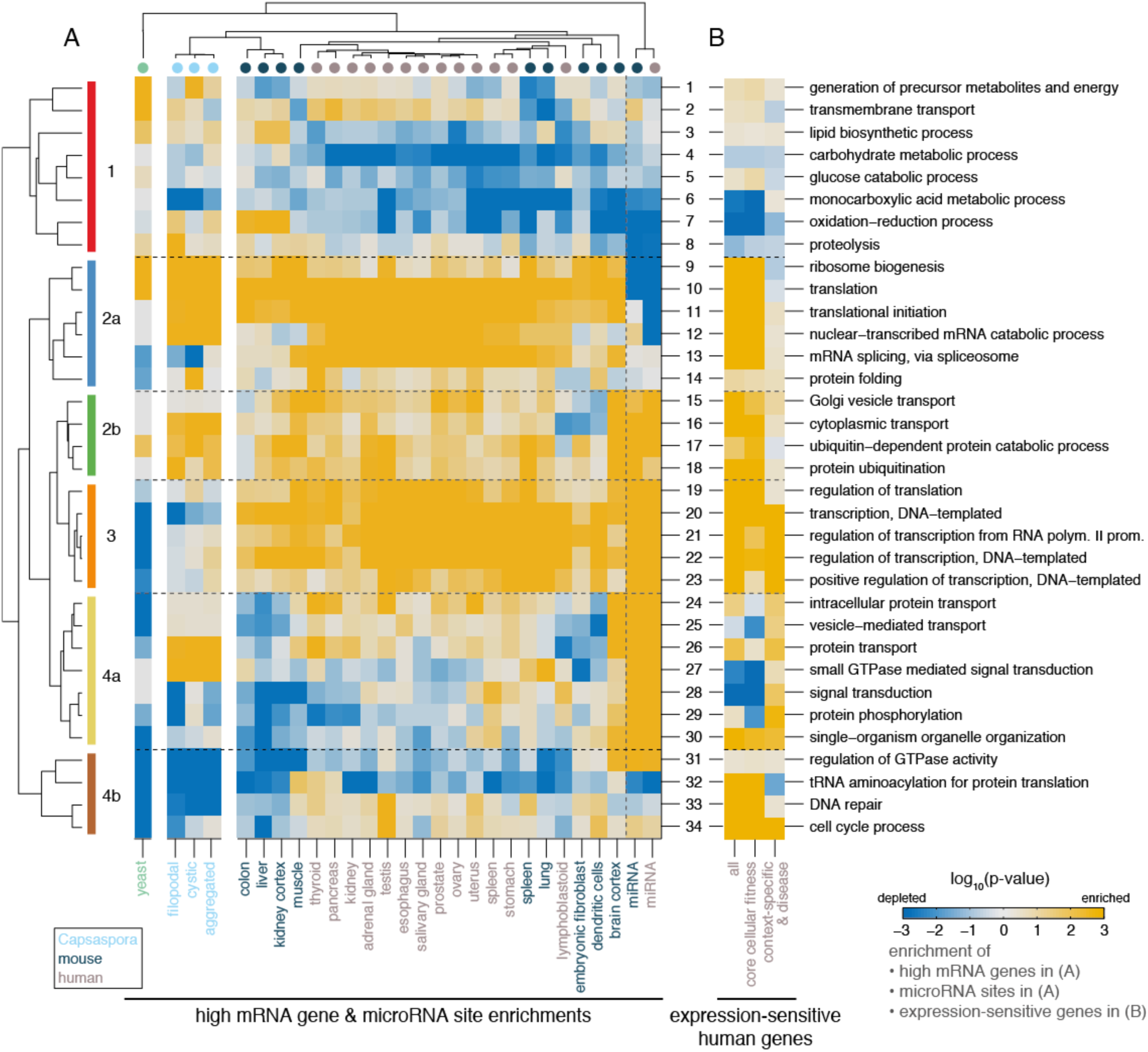
– Comparative analysis reveals evolutionary dynamics of mRNA/protein ratios. (A) Hierarchical clustering of high mRNA gene enrichment patterns in Gene Ontology biological processes (rows) across datasets (columns). Mouse and human datasets are shown individually with signed log_10_(p-values) for high mRNA gene enrichment. All yeast datasets as well as triplicate datasets of Capsaspora cell types were aggregated and combined p-values are shown (signed, log_10_). Human and mouse microRNA columns (right-most) are signed log_10_(p-values) for enrichment/depletion of microRNA binding sites. Biological processes are grouped into six (sub-)groups based on clustering (left side). (B) Heatmap of signed log_10_(p-values) of human expression-sensitive gene enrichment across GO biological processes, with rows ordered as in (A). Columns from left to right: all expression-sensitive genes, only core cellular fitness genes and only context-specific expression-sensitive genes (see Methods).

Hierarchical clustering of high mRNA gene and microRNA site enrichment patterns across biological processes reveals several groups of processes that show coherent similarities or differences across organisms (see Figure 5A and Table S1; see Figure 6A for organism-average enrichment effect sizes of processes; microRNA patterns are discussed below).

**Figure 6.**
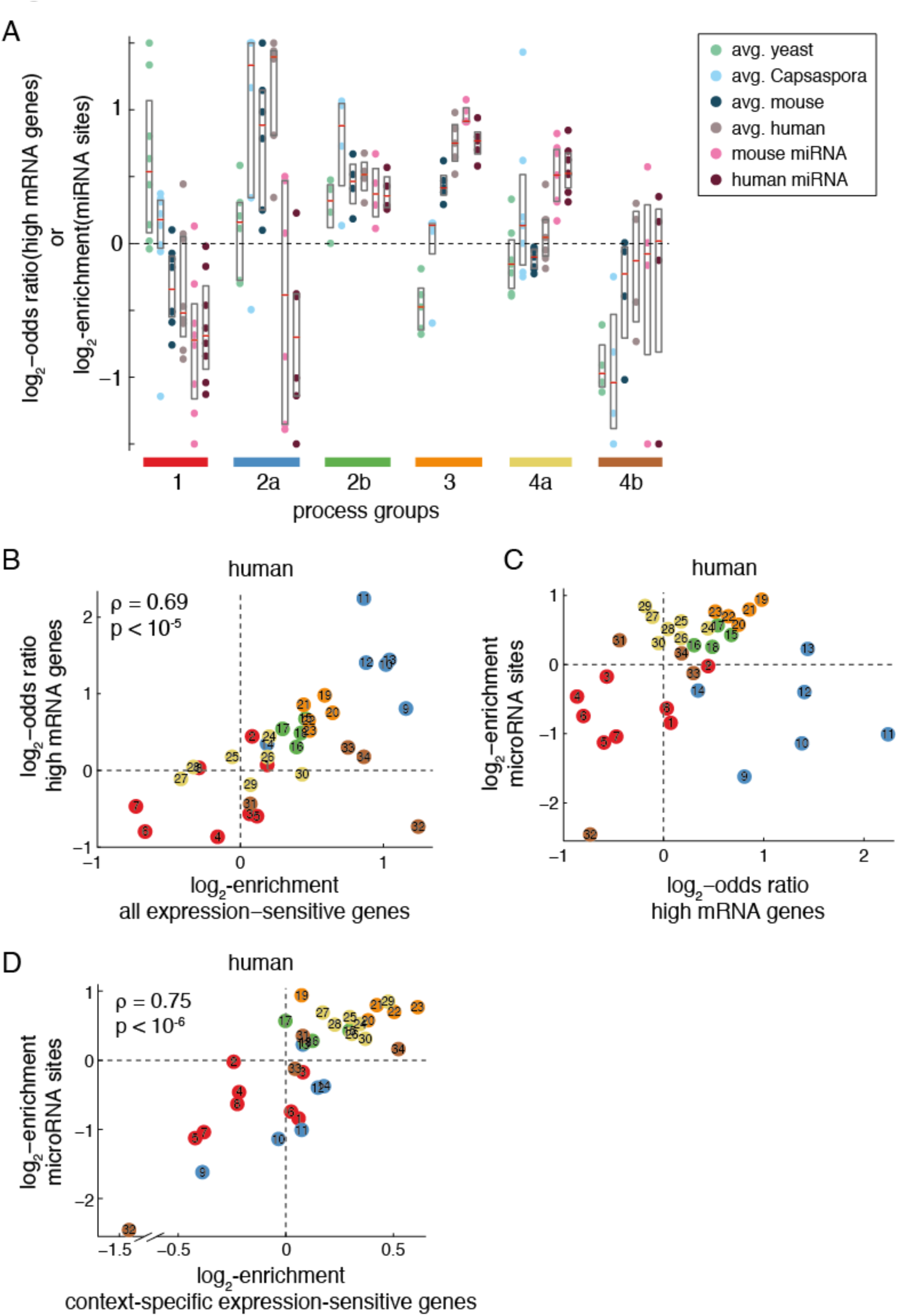
– Evolutionary dynamics of mRNA/protein ratios and their interdependency with microRNA regulation. (A) Species-average log_2_-odds ratios of high mRNA gene enrichment across biological process groups as defined in Figure 5A. Bars below the plot delineate groups. Shown for each group are the log_2_-odds ratios of high mRNA gene enrichment for each of its processes averaged across yeast, Capsaspora, mouse and human datasets, as well as the log_2_-enrichments for microRNA sites in mouse and human (from left to right). (B) Relationship between enrichment of human expression-sensitive genes and enrichment of high mRNA genes (human-average log_2_-odds ratios) across GO biological process terms. (C) Relationship between enrichment of high mRNA genes (human-average log_2_-odds ratios) and enrichment of conserved microRNA binding sites in human genes across GO biological processes. (D) Relationship between enrichment of human context-specific expressionsensitive genes and enrichment of conserved microRNA binding sites in human genes across GO biological processes. In (B-D), numbering and color-coding is according to Figure 5. Spearman’s rho and p-values are indicated in (B and D).

Metabolic processes (group 1) are enriched for high mRNA genes in yeast and mostly enriched in Capsaspora, but generally depleted of high mRNA genes in metazoan datasets, with the exception of mouse metabolic tissues.

Cellular core processes (group 2) are broadly enriched for high mRNA genes across all organisms. These include translation-related processes and protein folding (subgroup 2a) as well as protein ubiquitination and cytoplasmic transport processes (subgroup 2b).

To test whether this common enrichment of high mRNA genes reflects gene-specific conservation of mRNA/protein ratios, we investigated mRNA/protein ratios of orthologue pairs between human and each of the other three organisms. Orthologue pairs annotated for translation-related processes (processes 9-12) show significantly correlated mRNA/protein ratios, with correlation coefficients above those of all orthologues and increasing from yeast to Capsaspora to mouse (Spearman rank correlation, R = 0.31, p < 10^−3^, for human-yeast orthologues; R = 0.5, p < 10^−6^, for human-Capsaspora orthologues; R = 0.56, p < 10^−6^, for human-mouse orthologues; Supplementary Figure 7D). On the contrary, for genes annotated with subgroup 2b processes, only human-mouse orthologue pairs show an equivalently high correlation (Spearman rank correlation, R = 0.58, p < 10^−6^), but no significant correlation is found for human-yeast or human-Capsaspora orthologue pairs (Spearman rank correlation, R = 0.24, n.s.; R = -0.27, n.s.; respectively). Together this suggests that mRNA/protein ratios of genes in translation-related processes are subject to strong conservation across eukaryotes. On the other hand, subgroup 2b processes show a convergent tendency for large mRNA/protein ratio without detectable gene-specific conservation between unicellular eukaryotes and animals.

Regulatory processes (group 3), namely transcription, transcriptional regulation and translational regulation, are exclusively enriched for high mRNA genes in animal datasets. These processes are depleted for high mRNA genes in yeast, depleted/neutral in Capsaspora, but ubiquitously enriched for high mRNA genes in animal datasets, suggesting these are animal-specific gains of large mRNA/protein ratios.

Finally, a diverse set of processes involved in spatial organization, cell communication and surveillance (group 4) shows context-specific high mRNA gene enrichments in animal datasets. These processes are mostly depleted in yeast and show a range of enrichments in Capsaspora datasets but with little cell-type specific differences. In animal datasets, despite being on average across datasets mostly neutral, these processes show context-specific enrichments of high mRNA genes, such as protein transport and organelle organization in brain cortex, or signaling processes in immune-related tissues. Additionally, we find developmental and cell interaction processes to exhibit context-specific high mRNA gene enrichments in an animal-only analysis (Supplementary Figure 10 and Table S2).

In summary, post-transcriptional noise control has been lost during animal evolution for metabolic processes in most (non-metabolic) tissues, conserved or maintained at high levels for core cellular processes (group 2), gained across many cell-types for ubiquitously active regulatory processes (group 3) and gained in a context-specific manner for more specialized processes (group 4).

The state of high mRNA gene enrichments in processes from groups 1-3 is further supported by a positive correlation between average high mRNA gene enrichments and the fractions of expression-sensitive genes that biological processes contain (Spearman’s rho = 0.69, p < 10^−5^ in human; Figure 6B, see also Supplementary Figure 8 A&B). Furthermore, context-specific high mRNA gene enrichments of group 4a processes in animal datasets are supported by a high fraction of human genes with context-specific expression-sensitivities in these processes (Figure 5B). Together, this suggests that post-transcriptional noise control is adapted to the needs posed by evolving expression-sensitivities of genes.

### Animal microRNAs as evolutionary innovation for adaptive noise control

MicroRNA site enrichment patterns for biological processes in mouse and human are very similar to each other – and largely consistent with previous reports (Enright et al., 2003; John et al., 2004; Lewis et al., 2003; Stark et al., 2005) - but cluster separately from the enrichment of high mRNA genes in the four organisms (Figure 5A, see also Supplementary Figure 6B).

We find that the agreement between microRNA site enrichments and high mRNA gene enrichments depends on the process groups. For processes in groups 1, 2b and 3, enrichments of microRNA-binding sites correlate well with the average enrichments of high mRNA genes in human and mouse (Figure 6C and Supplementary Figures 8D and 11C). The data are consistent with microRNA sites being selected against where noise control has been lost (group 1) and selected for where noise control was maintained (group 2b) or gained (group 3) in animals. The previously observed enrichment of microRNA binding sites among genes with regulatory functions (Enright et al., 2003; John et al., 2004; Lewis et al., 2003; Stark et al., 2005) can thus be explained as a consequence of increasing needs for noise control during animal evolution.

Genes in group 2a processes are enriched for high mRNA genes but depleted of microRNA sites (Figure 6C). This depletion is in agreement with an enrichment for core cellular fitness genes (Figure 5B) and their shorter 3’UTRs, as well as microRNA site avoidance due to very large mRNA/protein ratios (Supplementary Figure 9). This therefore suggests that the depletion of microRNA binding sites among genes in cellular core processes results both from an avoidance of 3’UTR-mediated regulation (Stark et al., 2005) as well as from functional constraints in the noise control capacity of microRNAs.

Genes in processes with context-specific high mRNA gene enrichments in animal datasets (group 4a processes as well as developmental and cell communication processes from animal-only analysis, see Supplementary Figure 10 and Table S2) show strong enrichments of microRNA sites. These microRNA site enrichments, however, do not correlate with the average high mRNA gene enrichments of processes across datasets (Figure 6C and Supplementary Figure 11C). Instead, we find that they can be explained by a good agreement between microRNA site enrichments and the fractions of context-specific expression-sensitive genes per process (Spearman’s rho = 0.75, p < 10^−6^), which are high in those processes with context-specific high mRNA gene enrichments (Figure 6D and Supplementary Figures 8C and 11B). This is therefore in agreement with microRNAs controlling mRNA/protein ratios in genes whose expression-sensitivity depends on a specific cellular context.

In summary, high mRNA/protein ratios are a prevalent mechanism to control expression noise across a wide range of cellular processes. In contrast, microRNA/target interactions in animals evolved specifically for expression-sensitive genes in processes that require noise control in a specific cellular context in animals.

## Discussion

Fluctuations in protein numbers are an inevitable consequence of the stochasticity of chemical reactions. Our analyses show that increasing the number of mRNAs to produce proteins is a regulatory strategy to decrease protein fluctuations across organisms from distant parts of the tree of life. This suggests that noise control is a primary function of post-transcriptional mechanisms that alter protein expression level. Moreover, such a universal pattern substantiates the notion that gene expression noise is by its nature a detrimental force that all organisms have to manage.

Our results imply that post-transcriptional noise control could be a major reason why the genome-wide correlation between mRNA and protein levels is only moderate (Lawless et al., 2016; Schwanhäusser et al., 2011; Sebé-Pedrós et al., 2016b; Vogel et al., 2010; Wilhelm et al., 2014). While selection to minimize the metabolic cost of gene expression is expected to minimize the use of mRNAs (Frumkin et al., 2017; Wagner, 2005), selection to minimize noise posttranscriptionally will result in the increased use of mRNAs. The trade-off between these two opposing forces will thus lead to a range of gene-specific mRNA/protein ratios; with larger mRNA/protein ratios (i.e. lower translation rates) for those genes that organisms rely on the most.

Large mRNA/protein ratios are highly conserved for genes involved in translation and related processes across eukaryotes. However, our comparative analysis shows that post-transcriptional noise control is also adapted to organism-specific – and, for animals even to cell-type specific – requirements. Animals have developed a complex genomic architecture for transcriptional regulation (Sebé-Pedrós et al., 2016a) to establish developmental and cell-type specific expression programs (Arendt et al., 2016). Our data show that these regulatory processes acquired post-transcriptional noise control during animal evolution and suggest that the evolution of microRNAs played an important role in adapting noise control to the needs of specific expression programs. Together, these two evolutionary novel aspects of noise control might thus have facilitated the increasing organismal complexity of animals (Heimberg et al., 2008).

A scenario in which selection of microRNA-target interactions is chiefly driven by the need for noise control of expression-sensitive genes can help to explain why animal microRNA-target interactions – in contrast to those in plants (Jones-Rhoades et al., 2006) – individually mostly exert weak repressive effects (Baek et al., 2008; Lim et al., 2005; Selbach et al., 2008). That is, the need for precise expression levels attracts noise control but precludes large individual changes to expression levels. The establishment of combinatorial microRNA-target interactions and their respective compensatory increases in transcription would therefore have to proceed in gradual, alternating steps.

Our analyses prompt several questions. For instance, which are the mechanisms that control noise in translation and related processes, and are the mechanisms themselves conserved across eukaryotes? Furthermore, which other mechanisms are responsible for gains of noise control in animals, particularly for microRNA-independent gains in genes involved in mRNA splicing, cell cycle or immune processes. Further elucidating post-transcriptional noise control mechanisms may help to understand the plethora of post-transcriptional regulators as well as diseases that originate from mis-regulation at the post-transcriptional level. And finally, do transcriptional processes that affect expression noise exhibit similar bias towards expression-sensitive genes and how are they coordinated with post-transcriptional noise control?

Most importantly, here we have presented genome-wide evidence that microRNAs frequently function as noise controllers rather than as conventional repressors. Post-transcriptional noise control seems to be universal from bacteria to animals, with microRNAs specifically contributing to the evolution of noise control in regulatory and context-specific processes in animals.

## Acknowledgements

The authors would like to thank Arnau Sebé-Pedrós for sharing data and Nikolai Slavov for scientific discussion. This work was supported by the German Excellence Initiative (Humboldt Graduate School Post-Doc Scholarship, J.M.S.), the European Molecular Biology Organization (EMBO ALTF 857-2016, J.M.S.), the German Research Foundation (GRK 1772, N.B.) and the Berlin Institute of Health (TRG1, N.B.). Work in the lab of B.L. is funded by a European Research Council Consolidator grant (616434), the Spanish Ministry of Economy and Competitiveness (BFU2011-26206 and SEV-2012-0208), the AXA Research Fund, the Bettencourt Schueller Foundation, Agència de Gestió d’Ajuts Universitaris i de Recerca (AGAUR), the EMBL-CRG Systems Biology Program, and the CERCA Program/Generalitat de Catalunya.

## Author Contributions

Conceptualization, J.M.S., D.S.M., B.L. and N.B.; Methodology, J.M.S.; Formal Analysis, J.M.S.; Investigation, J.M.S.; Writing – Original Draft, J.M.S. and N.B.; Writing – Review & Editing, J.M.S., D.S.M, B.L. and N.B.; Supervision, D.S.M., B.L. and N.B.

## Materials & Methods

### Reference genomes

The following genomes and annotations were used for computations. *E.coli:* PEC (http://shigen.nig.ac.jp/ecoli/pec/about.jsp, June 2016), *S.cerviciae*: SGD (http://downloads.yeastgenome.org/curation/calculated_protein_info/, June 2016, build sacCer3). Capsaspora: *Capsaspora owczarzaki* genome v3. Mouse: RefSeq gene annotations build GRCm38/mm10 (Dec. 2011, downloaded from http://genome.ucsc.edu/). Human: RefSeq gene annotations build GRCh37/hg19 (Feb. 2009, downloaded from http://genome.ucsc.edu/)

### Gene sets

*E. coli* essential genes were obtained from PEC (http://shigen.nig.ac.jp/ecoli/pec/about.jsp, (Hashimoto et al., 2005)). Yeast haplo-insufficient genes were taken from Deutschbauer et al. (2005). Yeast genes sensitive to over-expression were taken from Makanae et al. (2013). Phenotypic effects of deletions in mice were obtained from the Mouse Genome Database consortium (Eppig et al., 2015) (http://www.informatics.jax.org/, November 2013) and essentiality of embryonic development was classified according to Georgi et al. (2013). Loss-of-function tolerant and recessive genes were taken from MacArthur et al. (2012). Haplo-insufficient disease genes (supported by PubMed or OMIM) were taken from Dang et al. (2008). Cancer genes were obtained from the Cancer Gene Census (Futreal et al., 2004) (http://cancer.sanger.ac.uk/census/), supplemented with 14 tumor suppressors and four oncogenes reported by Davoli et al. (2013). Enrichment for expression-altering mutations in cancer sequencing data were taken from Davoli et al. (2013) and genes classified as enriched for a specific mutation at q-values below 0.05. The set of genes reported as enriched in copy number amplifications was restricted to known oncogenes. Human genes essential for cell lines growth (growth phenotype) were obtained from Hart et al. (2015). One-to-one orthologues between human, mouse and yeast were downloaded from BioMart (Yates et al., 2016) (Ensembl 84). One-to-one orthologues between human and *Capsaspora owczarzaki* were kindly provided by Arnaud Sebe-Pedros (Sebé-Pedrós et al., 2016b).

### Analysis of microRNA regulation

Data on conserved microRNA binding sites in the 3’UTRs of mouse and human genes were downloaded from Targetscan (Garcia et al., 2011) (http://www.targetscan.org, version 6.2, June 2012). AU-rich elements [’ATTTA’, (Barreau et al., 2005)] and Pumilio 2 binding sites [’TGTANATA’, (Hafner et al., 2010)] in the 3’UTRs of mouse genes were determined using a custom Matlab script.

Expression parameters of genes in mouse embryonic fibroblasts (MEF) were obtained from Schwanhäusser et al. (2011) and an additional dataset of mRNA half-lives in MEF from Friedel et al. (2009). The number of conserved binding sites for microRNAs expressed in MEF [mouse microRNA expression atlas (Landgraf et al., 2007)] was calculated for all genes contained in the expression parameter dataset with a 3’UTR length of at least 100 nucleotides. Expression parameters of target sets of genes with varying numbers of conserved binding sites were compared to expression parameters of a 3’UTR length matched control set of genes without binding sites (‘background set’) and the relative difference of the medians was computed. Background sets with matched 3’UTR length were generated by selecting for each gene those ten genes with no conserved microRNA binding sites that are most similar in their 3’UTR length. Subsequently, the combined 3’UTR length distribution of target set and background set was spilt into ten equally spaced bins and the number of genes per bin between both distributions was matched by randomly removing surplus genes from the background set bins. By comparing the two sets using the Wilcoxon rank sum test on the length distribution, background sets that were significantly different to the target set at p<0.05 were discarded. The background set sampling procedure was repeated 1,000 times. The means and standard deviations of median ratios and the average p-value of all 1,000 iterations are reported.

Relation of expression parameters to the number of conserved binding sites for microRNAs expressed in MEF was also assessed by calculating the Spearman partial correlation coefficient between each expression parameter and the number of microRNA binding sites while controlling for the 3’UTR length of genes.

Relation of the total number of microRNA binding sites of genes in human (or mouse) to their mRNA/protein ratios across expression datasets was calculated as Spearman partial correlation coefficient between total number of conserved microRNA binding sites and mRNA/protein ratios in each expression dataset while controlling for 3’UTR length.

Enrichment or depletion of conserved microRNA binding sites of genes in a set compared to all genes not contained in the set was tested for significance with two-sided Wilcoxon rank sum test (Figure 1D-G).

### Expression and noise data sets

*E. coli.* Protein expression and protein noise data from YFP-fusion proteins and matched mRNA expression from RNA-sequencing were taken from Taniguchi et al. (2010). S. *cerevisiae*: Three datasets of protein expression measured by different mass spectrometry techniques were matched with mRNA expression data from RNA-sequencing (Lawless et al., 2016). An additional protein expression data set measured by mass spectrometry (Kulak et al., 2014) was combined with two mRNA expression datasets from RNA-sequencing (poly-A enrichment and RiboZero) (Presnyak et al., 2015). Protein expression and noise (split into total, intrinsic and extrinsic) measured from GFP-fusion proteins were obtained from Stewart-Ornstein et al. (2012). Two additional protein expression and total noise dataset from GFP-fusion proteins were obtained from Newman et al. (2006). *Capsaspora owczarzaki:* Triplicate datasets of mRNA (measured by RNA-sequencing) and protein (measured by mass spectrometry) expression in the three cell stages were obtained from Sebé-Pedrós et al. (2013) and Sebé-Pedrós et al. (2016b), respectively.

Mouse: Protein and mRNA expression data from embryonic fibroblasts were obtained from Schwanhäusser et al. (2011). Protein and mRNA expression data from bone marrow-derived dendritic cells were obtained from Jovanovic et al. (2015). For seven tissues (colon, kidney, liver, lung, skeletal muscle, spleen and brain cortex) mass spectrometry measurements of protein expression from Geiger et al. (2013) were matched to mRNA expression measured by RNA-sequencing from Merkin et al. (2012).

Human: Matched mRNA and protein expression data for twelve tissues (uterus, kidney, testis, pancreas, stomach, prostate, ovary, thyroid, adrenal gland, salivary gland, spleen, esophagus) were obtained from Wilhelm et al. (2014). An additional set of matched mRNA and protein expression data from a lymphoblastoid cell line was obtained from Khan et al. (2013).

For each dataset pair of matched mRNA and protein expression a running median of mRNA expression across the protein expression range was calculated with a window size of 5% of the number of genes contained in the datasets (but at least 50 genes). Genes with mRNA expression higher than the respective median were classified as *high mRNA genes*, others as *low mRNA genes*. Similarly, *high noise genes* and *low noise genes* were classified from comparison to running median of protein noise across protein expression range.

Fisher’s exact test was used to test enrichment or depletion of *high mRNA genes* (as well as *low noise genes*) in a gene set compared to all genes not contained in the set. Comparison of enrichment between two gene sets was assessed using Wilcoxon rank sum test. P-values from tests on multiple dataset pairs were combined using Fisher’s method.

### Comparison of microRNA targeting and mRNA/protein ratios

In each human transcriptome/proteome dataset, the distribution of log_10_-transformed mRNA/protein ratios was standardized (zero mean and unit standard deviation) and each gene’s standardized mRNA/protein ratios were averaged across datasets (‘scaled mRNA/protein ratios’).

Genes were grouped into bins with width 0.5 in scaled mRNA/protein ratio space, with the middle bin centered on 0. This bin size was chosen as a compromise between resolution and smoothness of resulting curves, but results are independent of bin size (data not shown). MicroRNA site density was calculated as the ratio of conserved microRNA binding sites and 3’UTR length, excluding genes with a 3’UTR shorter than 100 nucleotides.

The set *core cellular fitness genes* refers to cellular fitness genes that are essential in three or more out of five tested cell lines (Hart et al., 2015). The set *context*-*specific cellular fitness and disease genes* refers to the union of cellular fitness genes that are essential in less than three cell types and expression-sensitive disease genes (haplo-insufficient genes, tumor suppressors and oncogenes) that are not core cellular fitness genes.

### Gene Ontology enrichment analysis

Gene Ontology annotations for *S.cerevisiae* (GOC validation date 13/06/2016), mouse (GOC validation date 13/11/2014) and human (GOC validation date 21/03/2014) and the ontology (release 23/03/2014) were downloaded from http://geneontology.org/ (Ashburner et al., 2000; The Gene Ontology Consortium, 2015). Gene Ontology annotations for *Capsaspora owczarzaki* were kindly provided by Arnaud Sebé-Pedrós (Sebé-Pedrós et al., 2016b).

For each organism, only genes that are expressed in at least one of the matched mRNA and protein expression datasets were considered for further analysis (hereafter ‘expressed genes’). Biological process GO term annotations were mapped to expressed genes in the references genomes using the Matlab function goannotread. The list of gene-term associations was inflated by also mapping the respective one level-up parents of each term to annotated genes, in order to account for potential annotation differences between organisms. However, where the originally annotated child terms matched our applied thresholds for analysis (see below), superfluous parental terms were subsequently removed to minimize redundancy.

The threshold for consideration of a GO term in our analysis was a median of 30 annotated genes in the datasets of each species for all four species.

For each GO biological process, the log_2_-odds ratio of high mRNA gene enrichment in each dataset of matched mRNA and protein expression was calculated as described above and the significance of the enrichment/depletion was assessed using a one-sided Fisher’s exact test. Biological process enrichment for conserved microRNA binding sites was calculated as the average number of microRNA sites per expressed gene annotated for the process, divided by the average number of sites per expressed genes not annotated for the process. Significance of enrichment/depletion was tested using a Wilcoxon rank sum test.

Hierarchical clustering of high mRNA gene and microRNA site enrichment patterns was performed using cosine similarity of signed, log10-transformed p-values, i.e. *sign*(log_2_-odds) · |*log*_10_(p-value)|, of all individual datasets from yeast, Capsaspora, mouse and human. Because yeast as well as triplicate datasets from Capsaspora cell types show little intra-species/intra-cell type variation in GO term enrichment patterns, their datasets were subsequently summarized by calculating combined p-values of the datasets using Fisher’s method to improve clarity of the heat map.

Principal component analysis (Supplementary Figure 6) was performed using Matlab function *pca* on non-centered log_2_-odds ratios of high mRNA gene enrichments and log_2_-enrichments for microRNA sites.

Enrichment of GO terms with expression-sensitive genes was calculated on the set of expressed genes in each organism. Significance of enrichment/depletion was assessed using a one-sided Fisher’s exact test. For yeast, expression-sensitive genes are those determined as haplo-insufficient or overexpression-sensitive. In mouse, expression-sensitive genes are essential developmental genes, and enrichment was calculated based on the set of expressed genes that have also been tested for essentiality (Eppig et al., 2015). In human, *all expression*-*sensitive genes* refer to the set of genes encompassing cellular fitness genes (irrespective of number of cell types they are essential in), haplo-insufficient genes, tumor suppressors and oncogenes. As above, the set of *core cellular fitness genes* refers to cellular fitness genes that are essential in three or more out of five tested cell lines (Hart et al., 2015). The set of *context*-*specific cellular fitness and disease genes* refers to the union of cellular fitness genes that are essential in less than three cell types and expression-sensitive disease genes (haplo-insufficient genes, tumor suppressors and oncogenes) that are not core cellular fitness genes.

